# *In vitro* Evaluation of Antifungal Drug Combinations against Multidrug-resistant *Candida auris* isolates from New York Outbreak

**DOI:** 10.1101/593848

**Authors:** Brittany O’Brien, Sudha Chaturvedi, Vishnu Chaturvedi

## Abstract

Since 2016, New York hospitals and healthcare facilities have faced an unprecedented outbreak of pathogenic yeast *Candida auris*. We tested over one thousand *C. auris* isolates from affected facilities and found high-resistance to fluconazole (FLC, MIC_50_>256 mg/L), and variable resistance to other antifungal drugs. Therefore, we evaluated if two-drug combinations are effective *in vitro* against multidrug-resistant *C. auris*. Broth micro-dilution (BMD) plates were custom-designed, and quality controlled by TREK Diagnostic System. We used MIC_100_ endpoints for the drug combination readings as reported earlier for the intra- and inter-laboratory agreements against *Candida* species and *Aspergillus fumigatus*. The study results were derived from 12,960 MIC_100_ readings, for fifteen *C. auris* isolates tested against 864 possible two-drug antifungal combinations for nine antifungal drugs. Flucytosine (5FC) at 1.0 mg/L potentiated the most successful combinations with other drugs. Micafungin (MFG), Anidulafungin (AFG), Caspofungin (CAS) at individual concentrations of 0.25 mg/L in combination with 5FC (1.0 mg/L) yielded MIC_100_ for 14, 13, and 12 of 15 *C. auris* test isolates. AMB / 5FC (0.25/1.0 mg/L) yielded MIC_100_ for 13 isolates. None of the combinations were effective for *C. auris* 18-1, which tested resistant against FLC and 5FC, except POS/5FC (0.12/1.0 mg/L). The simplified two-drug combination susceptibility test format would permit laboratories to provide clinicians and public health experts with additional data to deal with multidrug-resistant *C. auris*.

## Introduction

*Candida auris* is an emerging multidrug-resistant yeast, causing invasive healthcare-associated infection with high mortality worldwide (1). It was described in 2009 as a new yeast species from the ear discharge of a Japanese patient (2). In the ensuing years, many reports of multidrug-resistant *C. auris* infections came from South Asia, South Africa, South America and travel-related cases from other parts of the world (3-6). The largest, localized, uninterrupted *C. auris* outbreak in the US healthcare facilities continues to afflict the New York metro areas (2013-2019); *C. auris* caused 23 deaths among the first 51 clinical case-patients in New York (7). In response, the New York State Department of Health (NYSDOH) laboratory scientists and epidemiologists developed enhanced protocols for the unprecedented surveillance and testing for *C. auris* (8). To date, over 15,000 clinical samples from 150 facilities were processed in the NYSDOH laboratory. The CLSI broth microdilution method (BMD) was used to test nearly one thousand *C. auris* isolates, which revealed high-resistance to fluconazole (MIC_50_ >256 mg/L), and variable resistance to other broad-spectrum antifungal drugs (Chaturvedi et al. unpublished data). Therefore, we investigated the efficacy of two-drug combinations against a representative set of fifteen drug-resistant *C. auris*.

## Materials and Methods

Isolates–*C. auris* isolates were initially processed on Sabouraud dulcitol agar at 40°C, and single colonies used to confirm identify with MALDI-TOF-MS and ITS -D1/D2 sequencing as described earlier. The isolates were stored at −80°C in 20% sterile glycerol. Drug combinations–Fluconazole (FLC) was not tested in combinations because of wide-spread resistance in our isolate collection. Nine drugs were selected for the combination testing: amphotericin B (AMB), anidulafungin (AFG), caspofungin (CAS), micafungin (MFG), flucytosine (5FC), isavuconazole (ISA), posaconazole (POS), voriconazole (VRC) and itraconazole (ITC). An advisory from the Centers for Diseases Control and Prevention was used for *C. auris* antifungal susceptibility testing and interpretation (https://www.cdc.gov/fungal/candida-auris/c-auris-antifungal.html). Combination testing–Broadly, we followed a method described earlier from a multi-laboratory evaluation of antifungal combination testing (9). Two fixed concentrations of each drug were tested, a value conforming to Cmax in the product insert, and a concentration considered resistant for a majority of pathogenic yeasts. The drug combinations were dispensed in a total volume of 100 μl. The 96-well microtiter plates were custom-manufactured and tested for sterility by TREK Diagnostic System, Part of Thermo Fisher Scientific, Oakwood Village, OH, USA. They were transferred under dry ice to NYSDOH Mycology Laboratory and stored at −80°Cuntil used. The schematic of plate design is provided in supplementary file 1. We also prepared two-fold dilutions of each drug combination by adding 100 μl sterile RPMI 1640 to the initial well and transferring 100 μl of 200 μl mix into micro-well of a new 96-well microtiter plate. This dilution scheme provided 1/2, 1/4, 1/8, 1/16, and 1/32 concentrations of each drug in the initial well. The inoculation, incubation, and MIC_100_ end-point reading, and *Candida krusei* ATCC was the quality control strain, were as published earlier from a multi-laboratory evaluation of antifungal combination testing (9). Briefly, *C. auris* cultures were streaked on Sabouraud dextrose agar for 24 hours at 35°C, cells suspended in sterile water to absorbance (A_530_) 0.08-0.1 absorbance as measured with a Mettler Toledo UV5 bio spectrophotometer; 20 μl of the cell suspension was added to the 11 ml of RPMI 1640 broth tube. One-hundred-microliter of the suspension in RPMI 1640 was added to each well of the 96-well microtiter plate with the drug combinations. The plates were incubated at 35°C for 48-hrs, and MIC_100_ reading was obtained with an illuminated convex mirror plate reader. Twenty microliters of the drug-fungal suspension from each well, after MIC_100_ determination, was transferred to the well of a fresh 96-well microtiter plate that contained 180 μl of sterile RPMI-1640 to set-up the sterility testing. We incubated the sterility plates at 35°C for 48-hrs, and determined the minimum fungicidal concentration (MFC) from each well with no visible growth. In preliminary experiments, a good correlation was observed between MFC values obtained from microbroth plates and Sabrouad dextrose agar (details not shown).

## Results

Among the test group, *C. auris* 16-1 was susceptible to all antifungals tested while the remaining fourteen isolates showed resistance to FLC (Table 1). The distribution of resistance in remaining *C. auris* was triazole resistance (2 isolates), echinocandin resistance (6 isolates), and AMB resistance (4 isolates) (Table 1). Only *C. auris* 18-1 and 18-13 showed resistance to 5FC. The combination test results were derived from 12,960 MIC_100_ readings, for fifteen *C. auris* isolates tested against 864 two-drug antifungal combinations for nine antifungal drugs (Supplementary Table 1). Flucytosine (5FC) at 1.0 mg/L potentiated the most successful combinations with other drugs (Table 2). MFG, AFG, and CAS at individual concentrations of 0.25 mg/L combined well with 5FC (1.0 mg/L) to yield MIC_100_ for 14, 13, and 12 out of 15 *C. auris* isolates tested. AMB / 5FC (0.25/1.0 mg/L) yielded MIC_100_ for 13 isolates. Triazoles POS, ISA and VRC also combined well with 5FC (0.25/2.0 mg/L) to yield MIC_100_ for 12, 13, and 13 isolates, respectively. The summary results for the remaining combinations are presented in supplementary table 2. Remarkably, MIC_100_ readings were obtained for all selected combinations. *C. auris* 18-1, resistant to FLC and 5FC initially, also tested resistant to all antifungal combinations except POS/5FC (0.12/1.0 mg/L) (Table 2).

**Table 1.** Broth microdilution determination of antifungal MIC_50_ (mg/L) and interpretative criteria (S, I, R) for the test isolates of *Candida auris*

| *C. auris* | Source | Antifungal drug |  |  |  |  |  |  |  |  |  |
| --- | --- | --- | --- | --- | --- | --- | --- | --- | --- | --- | --- |
| Susceptible |  | FLC | VOR | ITC | ISA | POS | AFG | CAS | MFG | AMB | 5FC |
| 16-1 | Ear | 8 | 0.25 | 0.5 | 0.25 | 0.25 | 0.12 | 0.12 | 0.12 | 0.5 | 0.5 |
|  |  | S | S | S | S | S | S | S | S | S | S |
| 17-12 | Urine | 32 | 0.5 | 0.06 | 0.015 | 0.03 | 8 | 8 | 8 | 1 | 0.032 |
|  |  | S | S | S | S | S | R | R | R | S | S |
| 18-14 | NA | >256 | 1 | 0.25 | 0.5 | 0.03 | 0.06 | 0.015 | 0.06 | 1 | 0.047 |
|  |  | R | S | S | S | S | S | S | S | S | S |
| 18-15 | Blood | >256 | 1 | 0.25 | 0.5 | 0.03 | 0.12 | 0.03 | 0.06 | 1.5 | 0.064 |
|  |  | R | S | S | S | S | S | S | S | S | S |
| 17-1 | Blood | >256 | 2 | 0.5 | 0.5 | 0.25 | 4 | 4 | 4 | 1 | 0.094 |
|  |  | R | S | S | S | S | R | R | R | S | S |
| 17-13 | Urine | >256 | 1 | 0.25 | 0.5 | 0.03 | 0.06 | 0.03 | 0.06 | 3 | 0.032 |
|  |  | R | S | S | S | S | S | S | S | R | S |
| 17-14 | Groin swab | >256 | 2 | 0.5 | 0.5 | 0.12 | 4 | 2 | 4 | 0.75 | 0.032 |
|  |  | R | S | S | S | S | R | R | R | S | S |
| 17-15 | Skin rash swab | >256 | 2 | 0.5 | 0.5 | 0.25 | 4 | 2 | 4 | 0.75 | 0.032 |
|  |  | R | S | S | S | S | R | R | R | S | S |
| 17-16 | Urine | >256 | 2 | 0.5 | 0.5 | 0.12 | 4 | 2 | 4 | 1 | 0.023 |
|  |  | R | S | S | S | S | R | R | R | S | S |
| 17-17 | Nares swab | >256 | 4 | 0.5 | 0.5 | 0.06 | 0.25 | 0.03 | 0.06 | 3 | 0.064 |
|  |  | R | R | S | S | S | S | S | S | R | S |
| 17-18 | Nares swab | >256 | 2 | 0.5 | 1 | 0.12 | 0.25 | 0.06 | 0.06 | 3 | 0.064 |
|  |  | R | S | S | S | S | S | S | S | R | S |
| 18-1 | Wound swab | 128 | 0.5 | 0.12 | 0.12 | 0.03 | 0.25 | 0.03 | 0.12 | 1 | >32 |
|  |  | R | S | S | S | S | S | S | S | S | R |
| 18-2 | Ascites fluid | >256 | 2 | 1 | 1 | 0.25 | 8 | 2 | 4 | 1.5 | 0.064 |
|  |  | R | S | R | R? | S | R | R | R | S | S |
| 18-5 | Urine | >256 | 2 | 0.25 | 0.5 | 0.06 | 0.12 | 0.03 | 0.12 | 2 | 0.094 |
|  |  | R | S | S | S | S | S | S | S | R | S |
| 18-13 | Rectal swab | 256 | 0.5 | 0.12 | 0.12 | 0.03 | 0.12 | 0.03 | 0.12 | 1 | >32 |
|  |  | R | S | S | S | S | S | S | S | S | R |

NA not available

**Table 2.** Most promising antifungal combinations for *Candida auris*

| Combi-<br>nation | End<br>-reading | *Candida auris* |  |  |  |  |  |  |  |  |  |  |  |  |  |  |
| --- | --- | --- | --- | --- | --- | --- | --- | --- | --- | --- | --- | --- | --- | --- | --- | --- |
|  |  | 16-1 | 17-1 | 17-12 | 17-13 | 17-14 | 17-15 | 17-16 | 17-17 | 17-18 | 18-1 | 18-2 | 18-5 | 18-13 | 18-14 | 18-15 |
| MFG +<br>5FC | MIC<sub>100</sub> | 0.12*/<br>1.0 | 0.12/<br>1.0 | 0.06/<br>0.5 | 0.12/<br>1.0 | 0.12/<br>1.0 | 0.12/<br>1.0 | 0.12/<br>1.0 | 0.12/<br>1.0 | 0.25/<br>2.0 | NA | 0.12/<br>1.0 | 0.12/<br>1.0 | 0.12/<br>1.0 | 0.12/<br>1.0 | 0.12/<br>1.0 |
| AMB +<br>5FC | MIC<sub>100</sub> | 0.25/<br>1.0 | 0.25/<br>1.0 | 0.25/<br>1.0 | 0.25/<br>1.0 | 0.25/<br>1.0 | 0.25/<br>1.0 | 0.25/<br>1.0 | 0.25/<br>1.0 | 0.25/<br>1.0 | NA | 0.25/<br>1.0 | 0.25/<br>1.0 | NA | 0.25/<br>1.0 | 0.25/<br>1.0 |
| AFG + 5FC | MIC<sub>100</sub> | 0.12 /<br>1.0 | 0.12/<br>1.0 | 0.12/<br>1.0 | 0.12/<br>1.0 | 0.12/<br>1.0 | 0.25/<br>2.0 | 0.12/<br>1.0 | 0.12/<br>1.0 | 0.25/<br>2.0 | NA | 0.12/<br>1.0 | 0.12/<br>1.0 | NA | 0.12/<br>1.0 | 0.12/<br>1.0 |
| CAS + 5FC | MIC<sub>100</sub> | 0.12/<br>1.0 | 0.12/<br>1.0 | 0.12/<br>1.0 | 0.12/<br>1.0 | 0.12/<br>1.0 | 0.12/<br>1.0 | 0.25/<br>2.0 | 0.12/<br>1.0 | NA | NA | 0.12/<br>1.0 | 0.25/<br>2.0 | NA | 0.12/<br>1.0 | 0.12/<br>1.0 |
| POS + 5FC | MIC<sub>100</sub> | 0.12/<br>1.0 | 0.12/<br>1.0 | 0.12/<br>1.0 | NA | NA | 0.12/<br>1.0 | 0.12/<br>1.0 | 0.12/<br>1.0 | 0.12/<br>1.0 | 0.12/<br>1.0 | 0.12/<br>1.0 | 0.12/<br>1.0 | NA | 0.12/<br>1.0 | 0.25/<br>2.0 |
| ISA + 5FC | MIC<sub>100</sub> | 0.12/<br>1.0 | 0.12/<br>1.0 | 0.12/<br>1.0 | 0.12/<br>1.0 | 0.12/<br>1.0 | 0.12/<br>1.0 | 0.12/<br>1.0 | 0.12/<br>1.0 | 0.12/<br>1.0 | NA | 0.12/<br>1.0 | 0.12/<br>1.0 | NA | 0.12/<br>1.0 | 0.25/<br>2.0 |
| ITC + 5FC | MIC<sub>100</sub> | 0.03/<br>0.5 | 0.06/<br>1.0 | 0.06/<br>1.0 | 0.06/<br>1.0 | NA | 0.06/<br>1.0 | 0.06/<br>1.0 | 0.06/<br>1.0 | 0.06/<br>1.0 | NA | 0.06/<br>1.0 | 0.06/<br>1.0 | NA | 0.06/<br>1.0 | 0.12/<br>2.0 |
| VOR + 5FC | MIC<sub>100</sub> | 0.25/<br>1.0 | 0.25/<br>1.0 | 0.125/<br>0.5 | 0.25/<br>1.0 | 0.25/<br>1.0 | 0.25/<br>1.0 | 0.25/<br>1.0 | 0.25/<br>1.0 | 0.25/<br>1.0 | NA | 0.25/<br>1.0 | 0.25/<br>1.0 | NA | 0.25/<br>1.0 | 0.5/<br>2.0 |

\* mg/L
NA Not available

## Discussion

The clinical and public health microbiology community is at high alert due to the alarming rise in the incidence and spread of drug-resistant *C. auris* globally (10). An aggressive therapeutic intervention is indicated for invasive *C. auris* infections associated with high mortality(7, 11). The results of the current study fill a gap in the published literature, as there is no report on the efficacy of antifungal combinations against *C. auris*. Our findings suggested that *C. auris* with various resistance patterns are susceptible to low dose combinations of existing drugs. Several of the effective combinations of antifungals exerted fungicidal activities, which was an additional indicator of the drug combination efficacy. The earlier multi-laboratory evaluation showed that MIC_100_ endpoints provide a reliable method for antifungal combination testing vis-a-vis ∑FIC (the summation of fractional inhibitory concentration) index (9). We further optimized the method by using fixed drug concentrations instead of the checkerboard dilutions, as described by other authors in earlier publications (12, 13). As almost all NY isolates belonged to the South Asian clade, it remains to be determined by other investigators if the resistant *C. auris* strains from South Africa and South America clades respond similarly to the antifungal combinations. Further clinical relevance of the findings reported in the study could be evaluated by pharmacokinetic/pharmacodynamic studies and testing in the animal models (14, 15). In conclusion, a limited set of two drug-combinations of antifungals were efficacious *in vitro* against fifteen multidrug-resistant *C. auris* strains.

The simplified two-drug combination susceptibility test format would permit laboratories to provide clinicians and public health experts with additional data to deal with multidrug-resistant *C. auris*.

## Supporting information

Supplement File 1- Combination plate design

Supplement File 2- Other effective combinations

## Acknowledgments

We are grateful to the members of the New York State Healthcare Epidemiology and Infection Control Program, *Candida auris* Investigation Workgroup, and the staff members from various hospitals and long-term-care facilities for assistance with surveillance samples.

## Transparency declaration

B. O’Brien performed the experiments and tabulated and analyzed the data, S. Chaturvedi supervised single drug testing and interpretation, helped interpreting combination testing and critiqued draft manuscript, and V. Chaturvedi conceived and designed the study, interpreted data, and wrote the manuscript.

## Supplementary Data

Supplementary File 1-Design of 96-well microtiter plates for the antifungal combination testing

Supplementary Table 1–Master file for all antifungal combination test results

Supplementary Table 2–Summary of less effective antifungal combinations against *Candida auris*

