## Supplement File 1- Combination plate design for "*In vitro* Evaluation of Antifungal Drug Combinations against Multidrug-resistant *Candida auris* isolates from New York Outbreak"

| | | | | | | | | |
|---|---|---|---|---|---|---|---|---|
| AMB 0.5 | FLC 2.0 | ITC 0.125 | AMB 0.5<br>FLC 2.0 | AMB 0.5<br>ITC 0.125 | AMB 2.0<br>FLC 2.0 | AMB 2.0<br>ITC 0.125 | AFG 0.25<br>ISA 0.25 | AFG 4.0<br>CAS 0.25 |
| AMB 2.0 | FLC 32.0 | ITC 1.0 | AMB 0.5<br>FLC 32.0 | AMB 0.5<br>ITC 1.0 | AMB 2.0<br>FLC 32.0 | AMB<br>2.0ITC 1.0 | AFG 0.25<br>ISA 2.0 | AFG 4.0<br>CAS 4.0 |
| AFG 0.25 | ISA 0.25 | AMB 0.5<br>AFG 0.25 | AMB 0.5<br>ISA 0.25 | AMB 2.0<br>AFG 0.25 | AMB 2.0<br>ISA 0.25 | AFG 0.25<br>CAS 0.25 | AFG 0.25<br>POS 0.25 | AFG 4.0<br>MFG 0.25 |
| AFG 4.0 | ISA 2.0 | AMB 0.5<br>AFG 4.0 | AMB 0.5<br>ISA 2.0 | AMB 2.0<br>AFG 4.0 | AMB 2.0<br>ISA 2.0 | AFG 0.25<br>CAS 4.0 | AFG 0.25<br>POS 1.0 | AFG 4.0<br>MFG 4.0 |
| CAS 0.25 | POS 0.25 | AMB 0.5<br>CAS 0.25 | AMB 0.5<br>POS 0.25 | AMB 2.0<br>CAS 0.25 | AMB 2.0<br>POS 0.25 | AFG 0.25<br>MFG 0.25 | AFG 0.25<br>VRC 0.5 | AFG 4.0<br>FLC 2.0 |
| CAS 4.0 | POS 1.0 | AMB 0.5<br>CAS 4.0 | AMB 0.5<br>POS 1.0 | AMB 2.0<br>CAS 4.0 | AMB 2.0<br>POS 1.0 | AFG 0.25<br>MFG 4.0 | AFG 0.25<br>VRC 2.0 | AFG 4.0<br>FLC 32.0 |
| MFG 0.25 | VRC 0.5 | AMB 0.5<br>MFG 0.25 | AMB 0.5<br>VRC 0.5 | AMB 2.0<br>MFG 0.25 | AMB 2.0<br>VRC 0.5 | AFG 0.25<br>FLC 2.0 | AFG 0.25<br>ITC 0.125 | AFG 4.0<br>ISA 0.25 |
| MFG 4.0 | VRC 2.0 | AMB 0.5<br>MFG 4.0 | AMB 0.5<br>VRC 2.0 | AMB 20<br>MFG 4.0 | AMB 2.0<br>VRC 2.0 | AFG 0.25<br>FLC 32.0 | AFG 0.25<br>ITC 1.0 | AFG 4.0<br>ISA 2.0 |

| | | |
|---|---|---|
| AFG 4.0<br>POS 0.25 | CAS 0.25<br>FLC 2.0 | **GC** |
| AFG 4.0<br>POS 1.0 | CAS 0.25<br>FLC 32.0 | CAS 0.25<br>ITC 0.125 |
| AFG 4.0<br>VRC 0.5 | CAS 0.25<br>ISA 0.25 | CAS 0.25<br>ITC 1.0 |
| AFG 4.0<br>VRC 2.0 | CAS 0.25<br>ISA 2.0 | CAS 4.0<br>MFG 0.25 |
| AFG 4.0<br>ITC 0.125 | CAS 0.25<br>POS 0.25 | CAS 4.0<br>MFG 4.0 |
| AFG 4.0<br>ITC 1.0 | CAS 0.25<br>POS 1.0 | CAS 4.0<br>FLC 2.0 |
| CAS 0.25<br>MFG 0.25 | CAS 0.25<br>VRC 0.5 | CAS 4.0<br>FLC 32.0 |
| CAS 0.25<br>MFG 4.0 | CAS 0.25<br>VRC 2.0 | **BL** |

| | | | | |
|---|---|---|---|---|
| CAS 4.0<br>ISA 0.25 | MFG 0.25<br>FLC 2.0 | MFG 0.25<br>ITC 0.125 | MFG 4.0<br>VRC 0.5 | FLC 2.0<br>VRC 0.5 |
| CAS 4.0<br>ISA 2.0 | MFG 0.25<br>FLC 32.0 | MFG 0.25<br>ITC 1.0 | MFG 4.0<br>VRC 2.0 | FLC 2.0<br>VRC 2.0 |
| CAS 4.0<br>POS 0.25 | MFG 0.25<br>ISA 0.25 | MFG 4.0<br>FLC 2.0 | MFG 4.0<br>ITC 0.125 | FLC 2.0<br>ITC 0.125 |
| CAS 4.0<br>POS 1.0 | MFG 0.25<br>ISA 2.0 | MFG 4.0<br>FLC 32.0 | MFG 4.0<br>ITC 1.0 | FLC 2.0<br>ITC 1.0 |
| CAS 4.0<br>VRC 0.5 | MFG 0.25<br>POS 0.25 | MFG 4.0<br>ISA 0.25 | FLC 2.0<br>ISA 0.25 | FLC 32.0<br>ISA 0.25 |
| CAS 4.0<br>VRC 2.0 | MFG 0.25<br>POS 1.0 | MFG 4.0<br>ISA 2.0 | FLC 2.0<br>ISA 2.0 | FLC 32.0<br>ISA 2.0 |
| CAS 4.0<br>ITC 0.125 | MFG 0.25<br>VRC 0.5 | MFG 4.0<br>POS 0.25 | FLC 2.0<br>POS 0.25 | FLC 32.0<br>POS 0.25 |
| CAS 4.0<br>ITC 1.0 | MFG 0.25<br>VRC 2.0 | MFG 4.0<br>POS 1.0 | FLC 2.0<br>POS 1.0 | FLC 32.0<br>POS 1.0 |

| | | | | | | |
|---|---|---|---|---|---|---|
| FLC 32.0<br>VRC 0.5 | ISA 0.25<br>ITC 0.125 | POS 0.25<br>VRC 0.5 | VRC 0.5<br>ITC 0.125 | | | **GC** |
| FLC 32.0<br>VRC 2.0 | ISA 0.25<br>ITC 1.0 | POS 0.25<br>VRC 2.0 | VRC 0.5<br>ITC 1.0 | | | |
| FLC 32.0<br>ITC 0.125 | ISA 2.0<br>POS 0.25 | POS 0.25<br>ITC 0.125 | VRC 2.0<br>ITC 0.125 | | | |
| FLC 32.0<br>ITC 1.0 | ISA 2.0<br>POS 1.0 | POS 0.25<br>ITC 1.0 | VRC 2.0<br>ITC 1.0 | | | |
| ISA 0.25<br>POS 0.25 | ISA 2.0<br>VRC 0.5 | POS 1.0<br>VRC 0.5 | | | | |
| ISA 0.25<br>POS 1.0 | ISA 2.0<br>VRC 2.0 | POS 1.0<br>VRC 2.0 | | | | |
| ISA 0.25<br>VRC 0.5 | ISA 2.0<br>ITC 0.125 | POS 1.0<br>ITC 0.125 | | | | |
| ISA 0.25<br>VRC 2.0 | ISA 2.0<br>ITC 1.0 | POS 1.0<br>ITC 1.0 | | | | **BL** |
