## Supplement File 2- Other effective combinations for "*In vitro* Evaluation of Antifungal Drug Combinations against Multidrug-resistant *Candida auris* isolates from New York Outbreak"

Supplementary Table 1. Other effective antifungal combinations against *Candida auris*

| Antifungal combinations | $MIC_{100}$ Concentration (No. positive isolates/ No. isolates tested) |
|---|---|
| MF + PZ | 0.125/0.125 (10/15) |
| AND + PZ | 0.125/0.125 (5/15) |
| CAS/PZ | 0.125/0.125 (5/15) |
| MF + ISA | 0.125/0.125 (5/15) |
| CAS + ISA | 0.125/0.125 (2/15) |
| AND + ISA | 0.125/0.125 (2/15) |
| MF + VOR | 0.125/0.25 (2/15) |
| AND + VOR | 0.0625/0.125 (1/15) |
| CAS + VOR | 0.125/0.25 (1/15) |
| MF + IZ | 0.125/0.06 (3/15) |
| CAS + IZ | 0.125/0.06 (2/15) |
| AND + IZ | 0.125/0.06 (2/15) |
| AB + MF | 0.5/0.25 (7/15) |
| AB + AND | 0.015/0.007 (1/15) |
| AB + CAS | 0.25/0.125 (1/15) |
| AB + PZ | 0.25/0.125 (5/15) |
| AB + ISA | 0.25/0.125 (2/15) |
| AB + IZ | 0.25/0.06 (2/15) |
| AB + VOR | 0.25/0.25 (1/15) |
| ISA + PZ | 0.125/0.125 (8/15) |
| PZ + VOR | 0.125/0.25 (7/15) |
| PZ + IZ | 0.125/0.06 (7/15) |
| ISA + IZ | 0.125/0.06 (2/15) |
| ISA + VOR | 0.125/0.25 (2/15) |
| VOR + IZ | 0.25/0.06 (2/15) |
| CAS + MF | 0.25/0.25 (2/15) |
| AND + MF | 0.125/0.125 (1/15) |
| AND/CAS | NA (0/15) |
