## Supplement File 3 for "*In vitro* Evaluation of Antifungal Drug Combinations against Multidrug-resistant *Candida auris* isolates from New York Outbreak"

| | | 16-1 | 17-1 | 17-12 | 17-13 |
|---|---|---|---|---|---|
| **Combination** | **Original concentration** | MIC100 | MIC100 | MIC100 | MIC100 |
| AB + AND | 0.5/0.25 | 0.25/0.125 | | | |
| AB + AND | 0.5/4 | 0.015/0.12 | 0.5/4 | 0.5/4 | 0.25/2 |
| AB + AND | 2/0.25 | 0.5/0.06 | | | |
| AB + AND | 2/4 | 0.06/0.12 | 1/2 | 2/4 | 0.5/1 |
| AB + CAS | 0.5/0.25 | 0.25/0.12 | | | |
| AB + CAS | 0.5/4 | 0.12/1 | | | |
| AB + CAS | 2/0.25 | 1/0.12 | 2/0.25 | | |
| AB + CAS | 2/4 | 0.25/0.5 | 2/4 | 1/2 | 2/4 |
| AB + MF | 0.5/0.25 | 0.25/0.12 | | | |
| AB + MF | 0.5/4 | 0.015/0.12 | | | 0.03/0.25 |
| AB + MF | 2/0.25 | 0.5/0.06 | | | 1/0.12 |
| AB + MF | 2/4 | 0.06/0.12 | 2/4 | | 0.12/0.25 |
| AB + FC | 0.5/2 | 0.25/1 | 0.25/1 | 0.25/1 | 0.25/1 |
| AB + FC | 0.5/32 | 0.03/2 | 0.015/1 | 0.015/1 | 0.03/2 |
| AB + FC | 2/2 | 0.5/0.5 | 0.5/0.5 | 0.5/0.5 | 1/1 |
| AB + FC | 2/32 | 0.06/1 | 0.06/1 | 0.06/1 | 0.12/2 |
| AB + ISA | 0.5/0.25 | 0.25/0.12 | | 0.25/0.12 | |
| AB + ISA | 0.5/2 | 0.03/0.12 | | 0.25/1 | |
| AB + ISA | 2/0.25 | 1/0.12 | | 1/0.12 | |
| AB + ISA | 2/2 | 0.06/0.06 | 2/2 | 0.5/0.5 | |
| AB + PZ | 0.5/0.25 | 0.25/0.12 | | 0.25/0.12 | |
| AB + PZ | 0.5/1 | 0.015/0.03 | 0.25/0.5 | 0.25/0.5 | 0.25/0.5 |
| AB + PZ | 2/0.25 | 0.5/0.06 | 1/0.12 | 1/0.12 | |
| AB + PZ | 2/1 | 0.06/0.03 | 1/0.5 | 0.5/0.25 | 1/0.5 |
| AB + VOR | 0.5/0.5 | 0.25/0.25 | | | |
| AB + VOR | 0.5/2 | 0.03/0.12 | | 0.25/1 | |
| AB + VOR | 2/0.5 | 1/0.25 | | | |
| AB + VOR | 2/2 | 0.25/0.25 | | 1/1 | |
| AB + IZ | 0.5/0.12 | 0.25/0.06 | | 0.25/0.06 | |
| AB + IZ | 0.5/1 | 0.25/0.5 | 0.25/0.5 | 0.25/0.5 | |
| AB + IZ | 2/0.12 | 1/0.06 | | 1/0.06 | |
| AB + IZ | 2/1 | 1/0.5 | 1/0.5 | 1/0.5 | |
| AND + CAS | 0.25/0.25 | | | | |
| AND + CAS | 0.25/4 | | | | |
| AND + CAS | 4/0.25 | 2/0.12 | | | 2/0.12 |
| AND + CAS | 4/4 | 1/1 | | | 4/4 |
| AND + MF | 0.25/0.25 | 0.12/0.12 | | | |
| AND + MF | 0.25/4 | 0.007/0.12 | | | 0.015/0.25 |
| AND + MF | 4/0.25 | 1/0.06 | | | 2/0.12 |
| AND + MF | 4/4 | 0.5/0.5 | | | 0.25/0.25 |
| AND + FC | 0.25/2 | 0.12/1 | 0.12/1 | 0.12/1 | 0.12/1 |
| AND + FC | 0.25/32 | 0.007/1 | 0.007/1 | 0.007/1 | 0.015/2 |
| AND + FC | 4/2 | 0.5/0.25 | 2/1 | 2/1 | 0.5/0.25 |
| AND + FC | 4/32 | 0.12/1 | 0.12/1 | 0.12/1 | 0.12/1 |
| AND + ISA | 0.25/0.25 | 0.12/0.12 | | 0.12/0.12 | |
| AND + ISA | 0.25/2 | 0.007/0.06 | | 0.12/1 | |

| | | | | | |
|---|---|---|---|---|---|
| AND + ISA | 4/0.25 | 0.12/0.007 | | 2/0.12 | 2/0.12 |
| AND + ISA | 4/2 | 0.12/0.06 | 4/2 | 2/1 | 1/0.5 |
| AND + PZ | 0.25/0.25 | 0.12/0.12 | | 0.12/0.12 | |
| AND + PZ | 0.25/1 | | | | |
| AND + PZ | 4/0.25 | 0.12/0.007 | | 2/0.12 | 2/0.12 |
| AND + PZ | 4/1 | 0.12/0.03 | 2/0.5 | 2/0.5 | 1/0.25 |
| AND + VOR | 0.25/0.5 | 0.06/0.12 | | | |
| AND + VOR | 0.25/2 | 0.015/0.12 | | 0.12/1 | |
| AND + VOR | 4/0.5 | 0.12/0.015 | | 2/0.25 | 2/0.25 |
| AND + VOR | 4/2 | 0.12/0.06 | | 2/1 | 2/1 |
| AND + IZ | 0.25/0.12 | 0.12/0.06 | | 0.12/0.06 | |
| AND + IZ | 0.25/1 | 0.06/0.25 | | 0.12/0.5 | 0.12/0.5 |
| AND + IZ | 4/0.12 | 0.5/0.015 | | 2/0.06 | 2/0.06 |
| AND + IZ | 4/1 | 0.12/0.03 | 2/0.5 | 2/0.5 | 2/0.5 |
| CAS + MF | 0.25/0.25 | 0.12/0.12 | | | |
| CAS + MF | 0.25/4 | 0.007/0.12 | | | 0.015/0.25 |
| CAS + MF | 4/0.25 | | | | |
| CAS + MF | 4/4 | 0.12/0.12 | | | |
| CAS + FC | 0.25/2 | 0.12/1 | 0.12/1 | 0.12/1 | 0.12/1 |
| CAS + FC | 0.25/32 | 0.015/2 | 0.007/1 | 0.007/1 | 0.015/2 |
| CAS + FC | 4/2 | 0.5/0.25 | 2/1 | 1/0.5 | 2/1 |
| CAS + FC | 4/32 | 0.12/1 | 0.12/1 | 0.12/1 | 0.5/4 |
| CAS + ISA | 0.25/0.25 | 0.12/0.12 | | 0.12/0.12 | |
| CAS + ISA | 0.25/2 | 0.007/0.06 | | 0.06/0.5 | |
| CAS + ISA | 4/0.25 | 0.25/0.015 | | 2/0.12 | |
| CAS + ISA | 4/2 | 0.12/0.06 | | 2/1 | 1/0.5 |
| CAS + PZ | 0.25/0.25 | 0.12/0.12 | | 0.12/0.12 | |
| CAS + PZ | 0.25/1 | 0.007/0.03 | 0.12/0.5 | 0.06/0.25 | 0.06/0.25 |
| CAS + PZ | 4/0.25 | 0.25/0.015 | 2/0.12 | 2/0.12 | 2/0.12 |
| CAS + PZ | 4/1 | 0.12/0.03 | 2/0.5 | 2/0.5 | 1/0.25 |
| CAS + VOR | 0.25/0.5 | 0.12/0.25 | | | |
| CAS + VOR | 0.25/2 | 0.015/0.12 | | 0.12/1 | |
| CAS + VOR | 4/0.5 | 0.5/0.06 | | 2/0.25 | |
| CAS + VOR | 4/2 | 0.25/0.12 | | 2/1 | 2/1 |
| CAS + IZ | 0.25/0.12 | 0.12/0.06 | | 0.12/0.06 | |
| CAS + IZ | 0.25/1 | 0.12/0.5 | | 0.12/0.5 | |
| CAS + IZ | 4/0.12 | 1/0.03 | | 2/0.06 | |
| CAS + IZ | 4/1 | | | | |
| MF + FC | 0.25/2 | 0.12/1 | 0.12/1 | 0.06/0.5 | 0.12/1 |
| MF + FC | 0.25/32 | 0.12/16 | | | 0.25/32 |
| MF + FC | 4/2 | 0.25/0.12 | 2/1 | 2/1 | 0.25/0.12 |
| MF + FC | 4/32 | 0.25/2 | | | 0.25/2 |
| MF + ISA | 0.25/0.25 | 0.06/0.06 | | 0.12/0.12 | |
| MF + ISA | 0.25/2 | 0.007/0.06 | | 0.12/1 | 0.12/1 |
| MF + ISA | 4/0.25 | 0.12/0.007 | | 2/0.12 | 0.5/0.03 |
| MF + ISA | 4/2 | 0.12/0.06 | | 0.25/0.12 | 0.5/0.25 |
| MF + PZ | 0.25/0.25 | 0.06/0.06 | 0.12/0.12 | 0.12/0.12 | 0.12/0.12 |
| MF + PZ | 0.25/1 | 0.015/0.06 | 0.12/0.5 | 0.015/0.06 | 0.12/0.5 |

| | | | | | |
|---|---|---|---|---|---|
| MF + PZ | 4/0.25 | 0.12/0.007 | | 2/0.12 | 0.25/0.015 |
| MF + PZ | 4/1 | 0.12/0.03 | 2/0.5 | 0.5/0.12 | 0.25/0.06 |
| MF + VOR | 0.25/0.5 | 0.06/0.12 | | | |
| MF + VOR | 0.25/2 | 0.015/0.12 | | 0.12/1 | 0.12/1 |
| MF + VOR | 4/0.5 | 0.12/0.015 | | | 0.25/0.03 |
| MF + VOR | 4/2 | 0.12/0.06 | | 2/1 | 0.25/0.12 |
| MF + IZ | 0.25/0.12 | 0.12/0.06 | | 0.12/0.06 | |
| MF + IZ | 0.25/1 | 0.06/0.25 | | 0.12/0.5 | |
| MF + IZ | 4/0.12 | 0.12/0.003 | | 2/0.06 | 0.25/0.0075 |
| MF + IZ | 4/1 | 0.12/0.03 | 2/0.5 | 2/0.5 | 0.25/0.06 |
| FC + ISA | 2/0.25 | 1/0.12 | 1/0.12 | 1/0.12 | 1/0.12 |
| FC + ISA | 2/2 | 0.06/0.06 | 1/1 | 0.25/0.25 | |
| FC + ISA | 32/0.25 | 2/0.015 | 1/0.007 | 1/0.007 | 2/0.015 |
| FC + ISA | 32/2 | 2/0.12 | 1/0.06 | 1/0.06 | 2/0.12 |
| FC + PZ | 2/0.25 | 1/0.12 | 1/0.12 | 1/0.12 | |
| FC + PZ | 2/1 | 0.06/0.03 | 1/0.5 | 0.12/0.06 | 1/0.5 |
| FC + PZ | 32/0.25 | 2/0.015 | 1/0.007 | 1/0.007 | 2/0.015 |
| FC + PZ | 32/1 | 1/0.03 | 1/0.03 | 1/0.03 | 2/0.06 |
| FC + VOR | 2/0.5 | 1/0.25 | 1/0.25 | 0.5/0.12 | 1/0.25 |
| FC + VOR | 2/2 | 0.12/0.12 | | 1/1 | |
| FC + VOR | 32/0.5 | 2/0.03 | 1/0.015 | 1/0.015 | 2/0.03 |
| FC + VOR | 32/2 | 4/0.25 | 1/0.06 | 1/0.06 | 2/0.12 |
| FC + IZ | 2/0.12 | 0.5/0.03 | 1/0.06 | 1/0.06 | 1/0.06 |
| FC + IZ | 2/1 | 1/0.5 | 1/0.5 | 1/0.5 | 1/0.5 |
| FC + IZ | 32/0.12 | 1/0.003 | 1/0.003 | 1/0.003 | 2/0.0075 |
| FC + IZ | 32/1 | 1/0.03 | 1/0.03 | 1/0.03 | 2/0.06 |
| ISA + PZ | 0.25/0.25 | 0.06/0.06 | 0.12/0.12 | 0.12/0.12 | |
| ISA + PZ | 0.25/1 | 0.007/0.03 | 0.12/0.5 | 0.015/0.06 | 0.12/0.5 |
| ISA + PZ | 2/0.25 | 0.06/0.007 | 1/0.12 | 0.25/0.03 | |
| ISA + PZ | 2/1 | 0.06/0.03 | | 0.25/0.12 | |
| ISA + VOR | 0.25/0.5 | 0.06/0.12 | | 0.12/0.25 | |
| ISA + VOR | 0.25/2 | 0.007/0.06 | 0.25/2 | 0.015/0.12 | 0.12/1 |
| ISA + VOR | 2/0.5 | 0.06/0.015 | | 0.25/0.06 | |
| ISA + VOR | 2/2 | 0.06/0.06 | | 0.12/0.12 | |
| ISA + IZ | 0.25/0.12 | 0.12/0.06 | | 0.12/0.06 | |
| ISA + IZ | 0.25/1 | 0.06/0.25 | 0.12/0.5 | 0.12/0.5 | |
| ISA + IZ | 2/0.12 | 0.06/0.003 | | 0.25/0.015 | |
| ISA + IZ | 2/1 | 0.06/0.03 | 1/0.5 | 0.12/0.06 | 1/0.5 |
| PZ + VOR | 0.25/0.5 | 0.06/0.12 | | 0.12/0.25 | |
| PZ + VOR | 0.25/2 | 0.12/1 | | 0.12/1 | |
| PZ + VOR | 1/0.5 | 0.03/0.015 | 0.5/0.25 | 0.12/0.06 | 0.5/0.25 |
| PZ + VOR | 1/2 | 0.03/0.06 | 0.5/1 | 0.06/0.12 | 0.5/1 |
| PZ + IZ | 0.25/0.12 | 0.06/0.03 | | 0.12/0.06 | |
| PZ + IZ | 0.25/1 | 0.06/0.25 | 0.12/0.5 | 0.06/0.25 | 0.12/0.5 |
| PZ + IZ | 1/0.12 | 0.03/0.003 | 0.5/0.06 | 0.12/0.015 | 0.5/0.06 |
| PZ + IZ | 1/1 | 0.03/0.03 | 0.5/0.5 | 0.06/0.06 | 0.5/0.5 |
| VOR + IZ | 0.5/0.12 | 0.25/0.06 | | 0.25/0.06 | |
| VOR + IZ | 0.5/1 | 0.12/0.25 | 0.25/0.5 | 0.25/0.5 | |

| | | | | |
|---|---|---|---|---|
| VOR + IZ | 2/0.12 | 0.25/0.03 | | 1/0.06 |
| VOR + IZ | 2/1 | 0.12/0.06 | 1/0.5 | 0.5/0.25 |

| 17-14 | 17-15 | 17-16 | 17-17 | 17-18 | 18-1 | 18-2 |
|---|---|---|---|---|---|---|
| MIC100 | MIC100 | MIC100 | MIC100 | MIC100 | MIC100 | MIC100 |
| | | | 0.06/0.5 | 0.12/1 | 0.06/0.5 | |
| | | | 1/0.12 | | 1/0.12 | |
| 2/4 | 2/4 | 2/4 | 0.25/0.5 | 0.5/1 | 0.25/0.5 | 2/4 |
| | | | | | 0.5/4 | |
| | | | 2/0.25 | | 2/0.25 | 2/0.25 |
| | 2/4 | 2/4 | 0.5/1 | 2/4 | 0.5/1 | 2/4 |
| | | | 0.25/0.12 | 0.5/0.25 | 0.25/0.12 | |
| | | | 0.015/0.12 | 0.03/0.25 | 0.015/0.12 | |
| | | | 1/0.12 | 2/0.25 | 1/0.12 | |
| | | | 0.06/0.12 | 0.12/0.25 | 0.06/0.12 | 2/4 |
| 0.25/1 | 0.25/1 | 0.25/1 | 0.25/1 | 0.25/1 | | 0.25/1 |
| 0.03/2 | 0.03/2 | 0.015/1 | 0.015/1 | 0.03/2 | | 0.015/1 |
| 1/1 | 1/1 | 1/1 | 1/1 | 1/1 | | 1/1 |
| 0.12/2 | 0.12/2 | 0.06/1 | 0.06/1 | 0.12/2 | | 0.06/1 |
| | | | | | 0.25/1 | |
| | | | | | 1/1 | |
| | | | | | 0.25/0.12 | 0.25/0.12 |
| 0.25/0.5 | 0.25/0.5 | 0.25/0.5 | 0.25/0.5 | 0.25/0.5 | 0.25/0.5 | 0.25/0.5 |
| | | | | | 1/0.12 | 1/0.12 |
| 0.5/0.25 | 1/0.5 | 0.5/0.25 | 1/0.5 | 1/0.5 | 0.5/0.25 | 0.5/0.25 |
| 0.25/0.5 | | 0.25/0.5 | 0.25/0.5 | | 0.25/0.5 | 0.25/0.5 |
| 1/0.5 | | 1/0.5 | 1/0.5 | | 1/0.5 | 1/0.5 |
| | | | 1/0.06 | 2/0.12 | 2/0.12 | |
| | | | 0.25/0.25 | 4/4 | 0.5/0.5 | |
| | | | 0.007/0.12 | 0.015/0.25 | 0.007/0.12 | |
| | | | 0.5/0.03 | 2/0.12 | 1/0.06 | |
| | | | 0.12/0.12 | 0.5/0.5 | 0.12/0.12 | |
| 0.12/1 | 0.25/2 | 0.12/1 | 0.12/1 | 0.25/2 | | 0.12/1 |
| 0.007/1 | 0.015/2 | 0.007/1 | 0.007/1 | 0.015/2 | | 0.007/1 |
| | 2/1 | 2/1 | 0.5/0.25 | 0.5/0.25 | 1/0.5 | 2/1 |
| 0.12/1 | 2/16 | 0.25/2 | 0.12/1 | 0.12/1 | 0.5/4 | 0.12/1 |
| | | | 0.12/1 | | 0.12/1 | |

| | | | | | | |
|---|---|---|---|---|---|---|
| | | | 2/0.12 | 2/0.12 | 0.5/0.03 | |
| 2/1 | 2/1 | 2/1 | 0.25/0.12 | 1/0.5 | 0.25/0.12 | 2/1 |
| | | | 0.12/0.12 | | 0.25/0.25 | 0.12/0.12 |
| | | | 2/0.12 | 2/0.12 | 2/0.12 | 2/0.12 |
| 2/0.5 | 2/0.5 | 2/0.5 | 0.25/0.06 | 1/0.25 | 0.5/0.12 | 2/0.5 |
| | | | | | 0.12/1 | |
| | | | 0.5/0.06 | 2/0.25 | 0.5/0.06 | |
| | | | 0.5/0.25 | 2/1 | 0.5/0.25 | |
| 0.12/0.5 | 0.12/0.5 | 0.12/0.5 | 0.12/0.5 | 0.12/0.5 | 0.12/0.5 | 0.12/0.5 |
| | | | 0.5/0.015 | 2/0.06 | 1/0.03 | |
| 2/0.5 | 0.25/0.06 | 2/0.5 | 1/0.25 | 2/0.5 | 1/0.25 | 2/0.5 |
| | | | 0.007/0.12 | 0.015/0.25 | 0.007/0.12 | |
| | | | 0.12/0.12 | | 0.25/0.25 | |
| 0.12/1 | 0.12/1 | 0.25/2 | 0.12/1 | | | 0.12/1 |
| 0.007/1 | 0.015/2 | 0.007/1 | 0.007/1 | 0.015/2 | | 0.007/1 |
| 2/1 | 2/1 | 2/1 | 0.5/0.25 | 1/0.5 | | 2/1 |
| 0.12/1 | 0.25/2 | 0.12/1 | 0.12/1 | 0.12/1 | | 0.12/1 |
| | | | | | 0.12/1 | |
| | | | 2/0.12 | | 2/0.12 | |
| 2/1 | | 2/1 | 1/0.5 | 1/0.5 | 1/0.5 | 2/1 |
| | | | | | 0.12/0.12 | 0.12/0.12 |
| 0.12/0.5 | 0.12/0.5 | 0.12/0.5 | 0.12/0.5 | 0.12/0.5 | 0.06/0.25 | 0.12/0.5 |
| 2/0.12 | 2/0.12 | 2/0.12 | 1/0.06 | 2/0.12 | 1/0.06 | 2/0.12 |
| 1/0.25 | 2/0.5 | 1/0.25 | 1/0.25 | 1/0.25 | 0.5/0.12 | 1/0.25 |
| | | | | | 2/0.25 | |
| | | | 1/0.5 | 2/1 | 1/0.5 | |
| 0.12/0.5 | 0.12/0.5 | | | | 0.12/0.5 | 0.12/0.5 |
| | | | 2/0.06 | | 2/0.06 | |
| 0.12/1 | 0.12/1 | 0.12/1 | 0.12/1 | 0.12/1 | | 0.12/1 |
| | | | 0.12/16 | | 0.25/32 | |
| 2/1 | 2/1 | 2/1 | 0.25/0.12 | 0.25/0.12 | 0.12/0.06 | 2/1 |
| | | | 0.12/1 | 0.25/2 | 0.12/1 | |
| | | | 0.12/0.12 | | 0.12/0.12 | |
| | | | 0.12/1 | | 0.12/1 | |
| | | | 0.12/0.007 | 0.25/0.015 | 0.12/0.007 | |
| | | | 0.12/0.06 | 0.25/0.12 | 0.12/0.06 | 2/1 |
| | 0.12/0.12 | 0.12/0.12 | 0.12/0.12 | | 0.12/0.12 | 0.12/0.12 |
| 0.12/0.5 | 0.12/0.5 | 0.06/0.25 | 0.12/0.5 | 0.12/0.5 | 0.06/0.25 | 0.12/0.5 |

| | | | | | | |
|---|---|---|---|---|---|---|
| | | | 0.12/0.007 | 0.12/0.007 | 0.12/0.007 | 2/0.12 |
| 2/0.5 | 2/0.5 | 2/0.5 | 0.12/0.03 | 0.25/0.06 | 0.12/0.03 | 2/0.5 |
| | | | 0.12/0.25 | | | |
| | | | 0.12/1 | | 0.12/1 | |
| | | | 0.12/0.015 | 0.25/0.03 | 0.12/0.015 | |
| | | | 0.12/0.06 | 0.25/0.12 | 0.12/0.06 | |
| | | | 0.25/0.12 | | | |
| | | 0.12/0.5 | 0.12/0.5 | | 0.12/0.5 | 0.12/0.5 |
| | | | 0.12/0.003 | 0.25/0.0075 | 0.12/0.003 | |
| 2/0.5 | | 2/0.5 | 0.12/0.03 | 0.25/0.06 | 0.12/0.03 | 2/0.5 |
| 1/0.12 | 1/0.12 | 1/0.12 | 1/0.12 | 1/0.12 | | 1/0.12 |
| | 1/1 | 1/1 | 1/1 | 1/1 | 1/1 | 1/1 |
| 1/0.007 | 2/0.015 | 1/0.007 | 1/0.007 | 1/0.007 | | 1/0.007 |
| 1/0.06 | 2/0.12 | 1/0.06 | 1/0.06 | 1/0.06 | 16/1 | 1/0.06 |
| | 1/0.12 | 1/0.12 | 1/0.12 | 1/0.12 | 1/0.12 | 1/0.12 |
| 1/0.5 | 1/0.5 | 0.5/0.25 | 1/0.5 | 1/0.5 | 0.5/0.25 | 1/0.5 |
| 1/0.007 | 2/0.015 | 1/0.007 | 1/0.007 | 1/0.007 | 16/0.12 | 1/0.007 |
| 1/0.03 | 2/0.06 | 1/0.03 | 1/0.03 | 1/0.03 | 8/0.25 | 1/0.03 |
| 1/0.25 | 1/0.25 | 1/0.25 | 1/0.25 | 1/0.25 | | 1/0.25 |
| | | | | | 1/1 | |
| 1/0.015 | 2/0.03 | 1/0.015 | 1/0.015 | 1/0.015 | | 1/0.015 |
| 1/0.06 | 2/0.12 | 1/0.06 | 1/0.06 | 1/0.06 | 16/1 | 1/0.06 |
| | 1/0.06 | 1/0.06 | 1/0.06 | 1/0.06 | | 1/0.06 |
| 1/0.5 | 1/0.5 | 1/0.5 | 1/0.5 | 1/0.5 | 1/0.5 | 1/0.5 |
| 1/0.003 | 2/0.0075 | 1/0.003 | 1/0.003 | 2/0.0075 | | 1/0.003 |
| 1/0.03 | 2/0.06 | 1/0.03 | 1/0.03 | 2/0.06 | 16/0.5 | 1/0.03 |
| | 0.12/0.12 | 0.12/0.12 | | | 0.12/0.12 | 0.12/0.12 |
| 0.06/0.25 | 0.12/0.5 | 0.12/0.5 | 0.12/0.5 | 0.12/0.5 | 0.06/0.25 | 0.12/0.5 |
| 1/0.12 | 1/0.12 | 1/0.12 | | | 1/0.12 | 1/0.12 |
| | | | | | 1/0.5 | |
| 0.06/0.5 | 0.12/1 | 0.12/1 | 0.12/1 | 0.12/1 | 0.03/0.25 | 0.06/0.5 |
| | | | | | 1/0.25 | |
| | | 1/1 | | | 1/1 | |
| 0.12/0.5 | 0.12/0.5 | 0.12/0.5 | | | 0.12/0.5 | 0.12/0.5 |
| | | | | | 1/0.06 | |
| 1/0.5 | 1/0.5 | 1/0.5 | 1/0.5 | 1/0.5 | 1/0.5 | 1/0.5 |
| | 0.12/0.25 | 0.12/0.25 | | | 0.12/0.25 | 0.12/0.25 |
| | 0.12/1 | 0.12/1 | | | 0.12/1 | 0.12/1 |
| 0.25/0.12 | 0.5/0.25 | 0.25/0.12 | | | 0.25/0.12 | 0.5/0.25 |
| 0.25/0.5 | 0.5/1 | 0.25/0.5 | 0.5/1 | 0.5/1 | 0.25/0.5 | 0.25/0.5 |
| | 0.12/0.06 | 0.12/0.06 | | | 0.12/0.06 | 0.12/0.06 |
| 0.12/0.5 | 0.12/0.5 | 0.12/0.5 | | 0.12/0.5 | 0.12/0.5 | 0.12/0.5 |
| 0.25/0.03 | 0.5/0.06 | 0.25/0.03 | 0.5/0.06 | 0.5/0.06 | 0.25/0.03 | 0.5/0.06 |
| 0.25/0.25 | 0.5/0.5 | 0.25/0.25 | 0.5/0.5 | 0.5/0.5 | 0.12/0.12 | 0.25/0.25 |
| 0.25/0.5 | 0.25/0.5 | 0.25/0.5 | | | 0.25/0.5 | 0.25/0.5 |

| | | | | | | |
|---|---|---|---|---|---|---|
| | | | | | 1/0.06 | |
| 1/0.5 | 1/0.5 | 1/0.5 | | | 1/0.5 | 1/0.5 |

| 18-5 | 18-13 | 18-14 | 18-15 |
|---|---|---|---|
| MIC100 | MIC100 | MIC100 | MIC100 |
| 0.5/4 | 0.06/0.5 | 0.5/4 | 0.5/4 |
| | 0.5/0.06 | 2/0.25 | 2/0.25 |
| 1/2 | 0.12/0.25 | 1/2 | 1/2 |
| | 0.5/4 | | |
| | 2/0.25 | | 2/0.25 |
| 2/4 | 0.5/1 | 2/4 | 2/4 |
| | 0.5/0.25 | 0.5/0.25 | 0.5/0.25 |
| 0.12/1 | 0.015/0.12 | 0.12/1 | 0.12/1 |
| 2/0.25 | 1/0.12 | 2/0.25 | 2/0.25 |
| 0.5/1 | 0.06/0.12 | 0.5/1 | 0.5/1 |
| 0.25/1 | | 0.25/1 | 0.25/1 |
| 0.015/1 | | 0.03/2 | 0.03/2 |
| 1/1 | 2/2 | 1/1 | 1/1 |
| 0.06/1 | | 0.12/2 | 0.12/2 |
| | 0.25/1 | | |
| | 1/1 | | |
| | 0.25/0.12 | | |
| 0.5/1 | 0.25/0.5 | 0.5/1 | 0.5/1 |
| | 1/0.12 | | |
| 2/1 | 1/0.5 | 1/0.5 | 2/1 |
| | 0.25/0.5 | | |
| | 1/0.5 | | |
| 4/0.25 | 1/0.06 | 4/0.25 | 4/0.25 |
| 4/4 | 0.5/0.5 | 4/4 | 4/4 |
| 0.25/4 | 0.007/0.12 | 0.06/1 | 0.06/1 |
| 4/0.25 | 0.5/0.03 | 4/0.25 | 4/0.25 |
| 4/4 | 0.12/0.12 | 2/2 | 2/2 |
| 0.12/1 | | 0.12/1 | 0.12/1 |
| 0.007/1 | | 0.015/2 | 0.015/2 |
| 1/0.5 | 0.5/0.25 | 1/0.5 | 1/0.5 |
| 0.12/1 | 0.5/4 | 0.12/1 | 0.12/1 |
| | 0.12/1 | | |

| | | | |
|---|---|---|---|
| 4/0.25 | 0.25/0.015 | 4/0.25 | 4/0.25 |
| 1/0.5 | 0.25/0.12 | 2/1 | 1/0.5 |
| | 0.12/0.12 | | |
| | 0.5/0.03 | 4/0.25 | 4/0.25 |
| 2/0.5 | 0.12/0.03 | 2/0.5 | 2/0.5 |
| | 0.12/1 | | |
| 4/0.5 | 0.25/0.03 | 4/0.5 | 4/0.5 |
| 4/2 | 0.25/0.12 | 4/2 | 2/1 |
| | 0.12/0.5 | | |
| 4/0.12 | 0.5/0.015 | 4/0.12 | 4/0.12 |
| 2/0.5 | 0.25/0.06 | 2/0.5 | 2/0.5 |
| | 0.25/0.25 | | |
| 0.06/1 | 0.015/0.25 | 0.06/1 | 0.06/1 |
| 0.25/2 | | 0.12/1 | 0.12/1 |
| 0.007/1 | | 0.015/2 | 0.015/2 |
| 2/1 | | 1/0.5 | 1/0.5 |
| 0.12/1 | | 0.12/1 | 0.25/2 |
| | 0.12/1 | | |
| | 2/0.12 | | 2/0.12 |
| 2/1 | 1/0.5 | 2/1 | 2/1 |
| | 0.12/0.12 | | |
| 0.25/1 | 0.06/0.25 | 0.25/1 | 0.25/1 |
| 2/0.12 | 1/0.06 | 2/0.12 | 2/0.12 |
| 2/0.5 | 1/0.25 | 4/1 | 2/0.5 |
| | 2/0.25 | | |
| 4/2 | 1/0.5 | 4/2 | 4/2 |
| | 0.12/0.5 | | |
| | 2/0.06 | | |
| 0.12/1 | 0.12/1 | 0.12/1 | 0.12/1 |
| | 0.12/16 | | |
| 1/0.5 | 0.12/0.06 | 0.5/0.25 | 1/0.5 |
| 2/16 | 0.12/1 | 1/8 | 1/8 |
| | 0.12/0.12 | | |
| | 0.12/1 | | |
| 4/0.25 | 0.12/0.007 | 2/0.12 | 1/0.06 |
| 1/0.5 | 0.12/0.06 | 1/0.5 | 1/0.5 |
| | 0.12/0.12 | | |
| 0.12/0.5 | 0.12/0.5 | 0.12/0.5 | 0.12/0.5 |

| | | | |
|---|---|---|---|
| 1/0.06 | 0.12/0.007 | 1/0.06 | 1/0.06 |
| 1/0.25 | 0.12/0.03 | 1/0.25 | 1/0.25 |
| | 0.12/1 | | |
| 4/0.5 | 0.12/0.015 | 1/0.12 | 1/0.12 |
| 2/1 | 0.12/0.06 | 1/0.5 | 1/0.5 |
| | 0.12/0.06 | | |
| | 0.12/0.5 | | |
| 2/0.06 | 0.12/0.003 | 1/0.03 | 1/0.03 |
| 2/0.5 | 0.12/0.03 | 1/0.25 | 1/0.25 |
| 1/0.12 | | 1/0.12 | 2/0.25 |
| 2/2 | | 1/1 | 2/2 |
| 1/0.007 | | 1/0.007 | 2/0.015 |
| 1/0.06 | 16/1 | 1/0.06 | 2/0.12 |
| 1/0.12 | 1/0.12 | 1/0.12 | 2/0.25 |
| 1/0.5 | 1/0.5 | 1/0.5 | 1/0.5 |
| 1/0.007 | 16/0.12 | 1/0.007 | 2/0.015 |
| 1/0.03 | 16/0.5 | 1/0.03 | 2/0.06 |
| 1/0.25 | | 1/0.25 | 2/0.5 |
| 1/0.015 | | 1/0.015 | 2/0.03 |
| 1/0.06 | | 1/0.06 | 2/0.12 |
| 1/0.06 | | 1/0.06 | 2/0.12 |
| 1/0.5 | 1/0.5 | 1/0.5 | 1/0.5 |
| 1/0.003 | | 1/0.003 | 2/0.0075 |
| 1/0.03 | 16/0.5 | 1/0.03 | 2/0.06 |
| | 0.12/0.12 | | |
| 0.25/1 | 0.12/0.5 | 0.12/0.5 | 0.12/0.5 |
| | 1/0.12 | | |
| | 1/0.5 | | |
| 0.25/2 | 0.12/1 | 0.12/1 | 0.12/1 |
| | 1/0.25 | | |
| | 1/1 | | |
| | 0.12/0.5 | | |
| | 1/0.06 | | |
| | 1/0.5 | | |
| | 0.12/0.25 | | |
| | 0.12/1 | | |
| 0.5/0.25 | 0.5/0.25 | 0.5/0.25 | 0.5/0.25 |
| 1/2 | 0.5/1 | 0.5/1 | 0.5/1 |
| | 0.12/0.06 | | |
| 0.25/1 | 0.12/0.5 | 0.12/0.5 | 0.12/0.5 |
| 1/0.12 | 0.5/0.06 | 0.5/0.06 | 0.5/0.06 |
| 0.5/0.5 | 0.5/0.5 | 0.5/0.5 | 0.5/0.5 |

| | | | |
|---|---|---|---|
| | 1/0.5 | | 2/1 |
